# Bleeding, cramping, and satisfaction among new copper IUD users: A prospective study

**DOI:** 10.1101/348094

**Authors:** Jessica N Sanders, Daniel E Adkins, Simranvir Kaur, Kathryn Storck, Lori M Gawron, David K Turok

## Abstract

**Objective:** We assess change in bleeding, cramping, and satisfaction among new copper (Cu) IUD users during the first six months of use, and evaluate the impact of bleeding and cramping on method satisfaction.

**Methods:** We recruited 77 women ages 18–45 for this prospective longitudinal observational cohort study. Eligible women reported regular menses, had no exposure to hormonal contraception in the last three months, and desired a Cu IUD for contraception. We collected data prospectively for 180 days following IUD insertion. Monthly, Participants reported bleeding scores using the validated pictorial blood loss assessment chart (PBAC), IUD satisfaction using a five-point Likert scale, and cramping using a seven-level ordinal scale. We used multiple imputation to address nonrandom attrition. Structural equation models for count and ordered outcomes modeled bleeding, cramping, and satisfaction growth curves over the six monthly repeated assessments.

**Results:** Bleeding significantly decreased (approximately 25%) over the course of the study from an estimated PBAC=195 at one month post-insertion to PBAC=151 at six months (t=−2.38, p<0.05). Additionally, IUD satisfaction improved over time (t=2.65, p<0.01), increasing from between “Neutral” and “Satisfied” to “Satisfied”, over the six month study. Cramping decreased sharply over the six-month study from between biweekly and weekly, to once or twice a month (t=−4.38, p<0.001). Finally, bleeding, but not cramping, was associated with IUD satisfaction (study mean: t=−2.31, p<0.05; study end: t=−2.81, p<0.01).

**Conclusions:** New Cu IUD users reported decreasing bleeding and cramping, and increasing IUD satisfaction, over the first six months. Method satisfaction was negatively associated with bleeding.

## Introduction

The intrauterine device (IUD) is the most common reversible method of contraception worldwide, with its use increasing in both developed and developing countries [1–3]. On average, 23% of contracepting women (with a range of <2% to >40%, depending on country) use either the hormonal or non-hormonal (copper) IUD [4]. At present, the copper (CuT380A) IUD is the only non-hormonal long-acting reversible contraceptive device approved by the United States Food and Drug Administration (FDA). The copper IUD is equally effective to sterilization and is designed for extended use (up to twelve years) [5]. However some women discontinue use due to undesired side effects. The most common reasons for discontinuation of a copper IUD during the first-year of use are symptoms related to pain or cramping and complaints of heavy bleeding [6, 7]. Perceived changes in vaginal bleeding may impact user preference, acceptability, method satisfaction and continuation [8].

There is a need for updated research evaluating IUD associated bleeding patterns for providers to better counsel patients on expectations when initiating a copper IUD. The primary objective of this study is to evaluate IUD associated bleeding and cramping patterns over the first six months after copper IUD insertion among individuals with regular menses who have not been exposed to any hormonal contraception in the past three months. A secondary objective of this study is to relate bleeding and cramping to IUD satisfaction in the months following insertion.

## Materials and methods

For this prospective cohort study, participants were recruited using convenience sampling through paper and electronic fliers, social media, and word-of-mouth, as well as by clinicians seeing patients for contraceptive services. A trained research assistant screened individuals interested in the copper IUD for contraception over the phone to ensure they met study criteria. Women, aged 18-45 years, with self-reported normal menses, lasting between three and seven days, were eligible for participation. Eligible participants needed to be able to provide bleeding information for three cycles free of any hormonal contraception prior to enrollment and be willing to track bleeding for an additional 180 days. Individuals who were six or more months postpartum, and not breastfeeding, were eligible if they had returned to normal menses for three cycles.

Exclusion criteria included use of any hormonal contraceptive methods in the three months, irregular length of any of the previous three menstrual cycles (less than 21 days, or longer than 35 days), or irregular duration of vaginal bleeding in the previous three cycles (less than three days, or longer than seven days) at the time of screening. Other exclusion criteria included any medical contraindication to the copper T380A IUD per the package insert [9], desire for pregnancy within six months, and participation in any other clinical research investigation.

This study aimed to incorporate validated methods to evaluate bleeding profiles with electronic data capture and popular menstrual tracking mobile applications to examine changes in bleeding patterns among women initiating the copper IUD [10–12]. The analysis uses two validated measures: the World Health Organization (WHO) recommendations and the pictorial blood loss assessment chart (PBAC). The WHO method includes data collection on bleeding days (defined as days when blood loss requires the use of a menstrual pad or tampon) and spotting days (defined as bloody vaginal discharge, but when no protection was needed) [10]. The second tool, the PBAC, is an inexpensive and reliable method of assessing frequency, duration, and intensity of vaginal bleeding [12]. We calculated monthly PBAC scores based on the published recommendations: one point for each lightly stained pad or tampon reported, five points for moderately saturated pads or tampon, and 10 points for a completely saturated tampon or 20 points for completely saturated pad. Additional points were added for small clots (1pt), large clots (5pt), or flooding (5pt). We also assessed the frequency of the following hormonal/menstrual symptoms adapted from the Menstrual Symptoms Questionnaire (MSQ): headache, bloating, breast tenderness, moodiness/irritability, acne, cramping, weight gain, weight loss, depression, constipation and, diarrhea [13]. Additionally, participants completed a monthly survey that assessed IUD use, satisfaction, and potential adverse events. We assessed contraceptive method satisfaction using a five-point Likert scale (very unsatisfied, unsatisfied, neutral, satisfied, very satisfied) and cramping frequency was assessed using a seven-level ordinal scale (never to every day of the month).

We used the Research Electronic Data Capture system (REDcap), a secure online database for survey creation, data collection and management,[14] to collect survey data and bleeding diaries. Retrospective data were collected for the 90 days prior to copper IUD insertion and prospective data was collected for 180 days after insertion using REDCap. We conducted repeated measures analysis at nine time points: three months before insertion and monthly for six months after insertion. Participants were encouraged to use electronic period trackers or daily diaries to track the information that they entered monthly into the REDCap survey.

We modeled longitudinal growth curve trajectories of bleeding scores, IUD satisfaction, and cramping (as reported by the MSQ) using structural equation models (SEM) for count and ordinal outcomes.[15–17] This modeling strategy permits the specification of fixed and random intercepts and slopes allowing us to formally test whether bleeding decreases, on average, across the study course (fixed time effect), and whether the rate of change varies across individuals (random time effect).

As none of the three outcomes are continuous variables (instead, they are a count variable [i.e., PBAC score assessing bleeding] and two ordinal variables [i.e., IUD satisfaction Likert scale; cramping MSQ item]), we use models for discrete outcomes. Specifically, we use count link functions for the bleeding growth curve, and ordinal link function for the satisfaction and cramping growth curves [17, 18]. Given marked overdispersion in the bleeding measure (mean=148.4; variance=21242.2), we present negative binomial results in the main text and verify the robustness using Poisson and standard linear link functions in the Supporting Information. For the ordinal measures, IUD satisfaction and cramping, we present the conventional ordered logistic link function results in the main text and verify the robustness of the specification using ordered probit and linear link functions.

We anticipated a missing data pattern in which women with less favorable outcomes are more likely to attrite. To avoid biased estimates overstating bleeding and cramping improvements over the study period we used a well-validated multiple imputation (MI) method (i.e., multiple imputation with chained equations [MICE}) [19]. We used a conservative 30 imputations in all MI analyses, which were conducted using the “mi estimate” command in Stata. Finally, we output subject-level estimates for post-insertion bleeding and post-insertion cramping from the growth curve models, as well as retrospectively reported baseline bleeding, and tested their association to IUD method satisfaction (specified as both mean satisfaction across the study, and satisfaction at study conclusion).

We informed the power calculations from two prior studies [11, 20]. Belsey found that women on non-hormonal methods had a mean length of bleeding/spotting lasting five days out of 30 [21] and Tolley et al. found that each additional day of bleeding increased the likelihood of discontinuation by about 3% for IUD users [20]. Thus, we estimated that changes longer than 2 days may be considered a menstrual disruption and could potentially impact user satisfaction and continuation. Using these assumptions with a standard deviation of 4 days, a sample size of 64 would allow for 80% power with an alpha of 0.05 to detect 2-day change in bleeding. Our enrollment goal of 77 accounted for a 20% loss to follow up. This study protocol was submitted to ClinicalTrials.gov (Identifier NCT02311478). The University of Utah Institutional Review Board approved this study. Teva Pharmaceutical Industry, Ltd. funded this investigator initiated study and reviewed the manuscript prior to submission. All analyses were conducted in Stata 15.1.

## Results and discussion

We enrolled 79 women into this study, 78 women had a successful IUD insertion. One individual immediately withdrew from the study and 5 (8%) were lost to follow-up after the insertion. Seventy-two women (92%) completed the 6-month follow up: 1 (0.01%) participant had an expulsion, 8 (11%) elected to have removals, and 63 (88%) participants were still using their device at 6 months (Supporting Figure S1). The mean age of participants was 25 years (SD 5.2). Three-quarters of women reported using hormonal contraception in the past, with a median of nine months (mean 18.7 months; SD 25.3 range 3-120 months) since last use. Most participants reported being single/non-cohabitating (62%) and had a normal body mass index (18.5 - 24.9kg/m^2^) (65%). In the enrollment survey, half (49%) of women reported 3 to 4 days of bleeding per month and half (51%) reported 5 to 7 days of bleeding per month. Over the course of the study the total accumulated person-months for key assessments ranged between 406 (88%) for IUD satisfaction and 415 (90%) for bleeding outcomes. In all MI analyses, there were 462 person months (77 subjects*6 assessments). For additional description of participants see Table 1. The MI distributions for longitudinal outcomes of bleeding, cramping and IUD satisfaction are depicted in the dotplots in Figure 1 A-C.

**Table 1.**
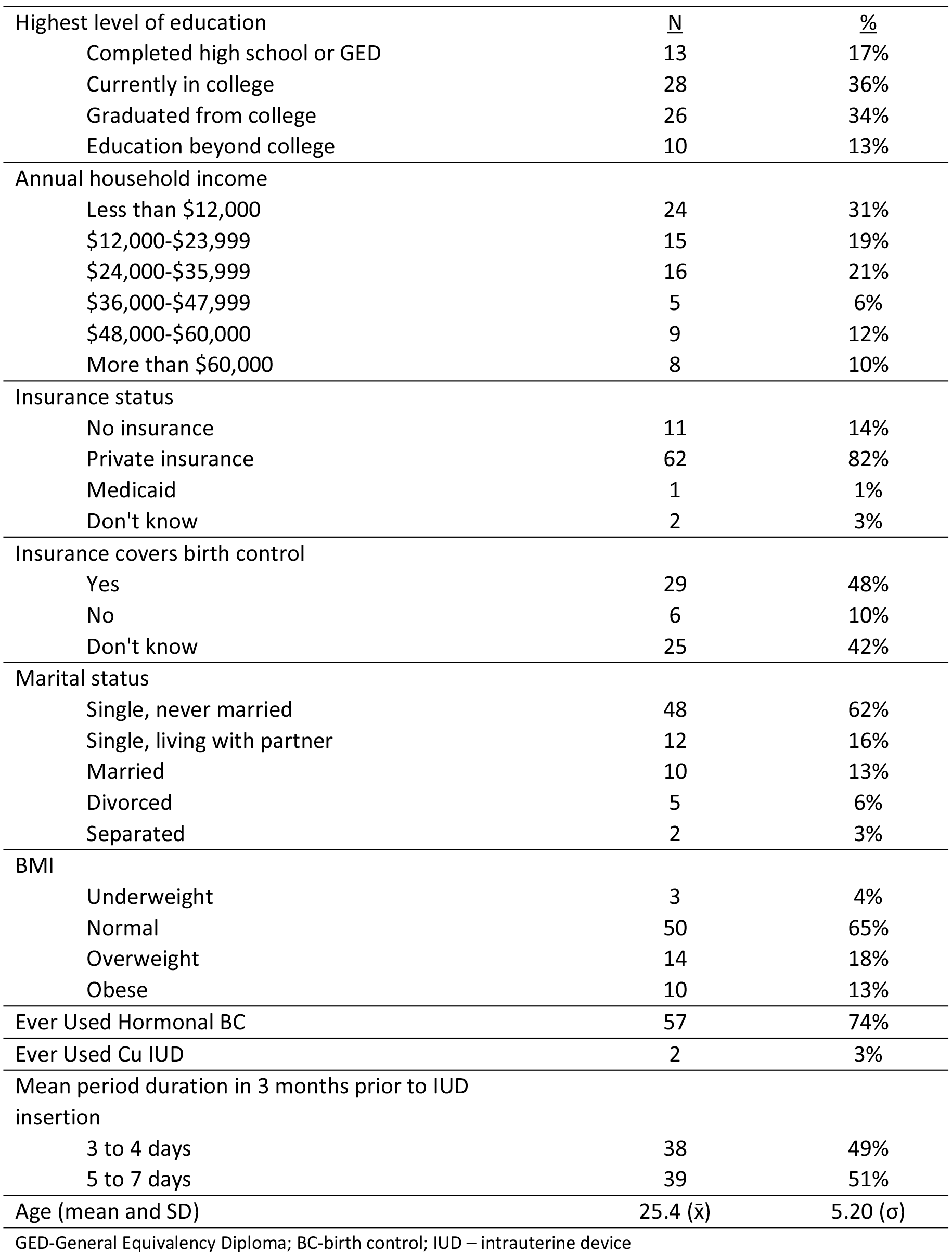
Sample summary statistics (N=77)

| Highest level of education | <u>N</u> | <u>%</u> |
| --- | --- | --- |
| Completed high school or GED | 13 | 17% |
| Currently in college | 28 | 36% |
| Graduated from college | 26 | 34% |
| Education beyond college | 10 | 13% |
| Annual household income |  |  |
| Less than \$12,000 | 24 | 31% |
| \$12,000-\$23,999 | 15 | 19% |
| \$24,000-\$35,999 | 16 | 21% |
| \$36,000-\$47,999 | 5 | 6% |
| \$48,000-\$60,000 | 9 | 12% |
| More than \$60,000 | 8 | 10% |
| Insurance status |  |  |
| No insurance | 11 | 14% |
| Private insurance | 62 | 82% |
| Medicaid | 1 | 1% |
| Don't know | 2 | 3% |
| Insurance covers birth control |  |  |
| Yes | 29 | 48% |
| No | 6 | 10% |
| Don't know | 25 | 42% |
| Marital status |  |  |
| Single, never married | 48 | 62% |
| Single, living with partner | 12 | 16% |
| Married | 10 | 13% |
| Divorced | 5 | 6% |
| Separated | 2 | 3% |
| BMI |  |  |
| Underweight | 3 | 4% |
| Normal | 50 | 65% |
| Overweight | 14 | 18% |
| Obese | 10 | 13% |
| Ever Used Hormonal BC | 57 | 74% |
| Ever Used Cu IUD | 2 | 3% |
| Mean period duration in 3 months prior to IUD insertion |  |  |
| 3 to 4 days | 38 | 49% |
| 5 to 7 days | 39 | 51% |
| Age (mean and SD) | 25.4 ($\bar{x}$) | 5.20 ($\sigma$) |

GED-General Equivalency Diploma; BC-birth control; IUD – intrauterine device

**Fig 1.**
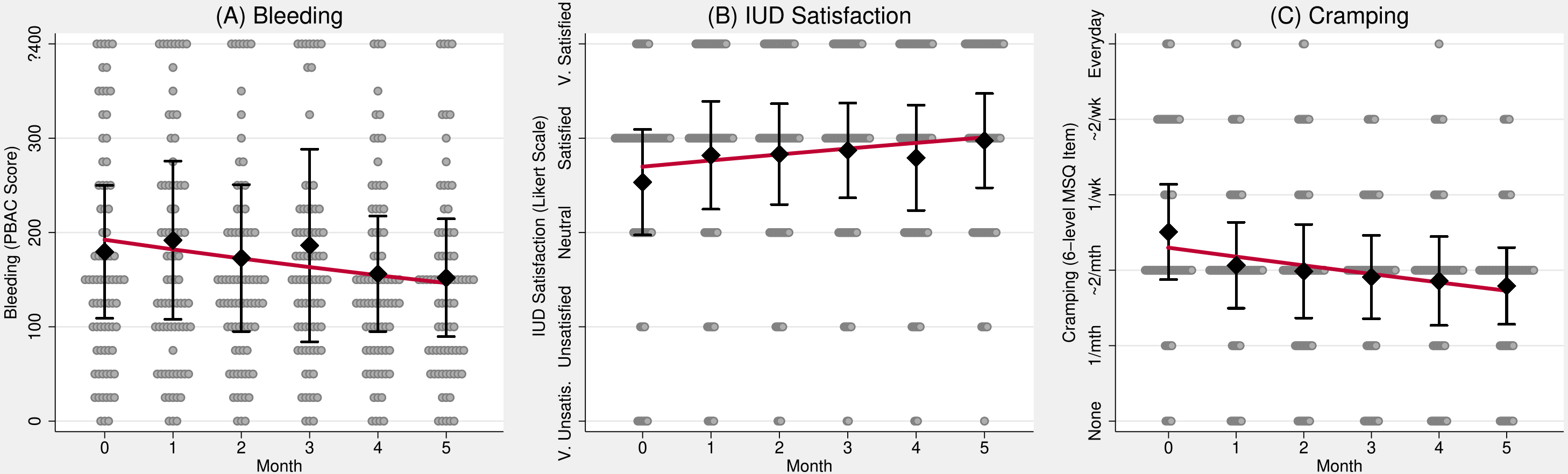
Time series plots summarizing distributions, and predicted mean growth curves, of (A) Bleeding, (B) IUD Satisfaction, and (C) Cramping over study month. Black diamonds represent variable means (i.e., Bleeding, IUD satisfaction, and Cramping), by month. Interval bars represent variable means +/− 0.5 variable SDs, by month. The red lines represent the model-implied growth curve as predicted by the SEM models presented in Table 2. The underlying dotplot depicts the distributions of the outcomes, by month. For the first plot, (A), values are binned, and the Y axis is top coded at ≥400 for the PBAC score to improve resolution in the middle of the distribution. Abbreviations: PBAC – Pictorial Blood Assessment Chart; IUD – intrauterine device; MSQ - Menstrual Symptoms Questionnaire

As shown in the second column of Table 2, the fixed time slope coefficient for the bleeding negative binomial SEM growth curve was significantly negative (b=−0.055, p<0.05), indicating that bleeding declined over the six-month study period. Comparison of the complete case analysis (see Supplemental Table 1) to the MI findings of the same outcome (Supplemental Table 2) indicate that the MI approach captures and corrects a moderate attrition bias; and contributes robust evidence for significant decrease in bleeding in the six-month period following IUD insertion. As visualized in Figure 1, Panel A, the significant random intercept term (p<0.001) indicates significant variation in baseline bleeding between women in the study, and our growth curve estimates (visualized as a red line) indicate a ˜25% reduction in bleeding across the six-month study period. Cramping also varied across women at at baseline, and notably decreased from between bimonthly and weekly to once or twice a month across the six-month study period (b=−1.77; p<0.001; Figure 1, panel B).

The IUD satisfaction ordered logistic SEM growth curve (Figure 1, panel C) indicated a significant increase in IUD satisfaction in the six months following placement (b=0.154; p<0.01). As depicted in Figure 1 Panel B, the model-implied IUD satisfaction growth curve increases about one third of a Likert category from between “Neutral” and “Satisfied” to “Satisfied”, on average, across the six months following IUD placement. Comparing IUD satisfaction complete case results (Supplemental Table 3) to MI results (Supplemental Table 4), and cramping complete case results (Supplemental Table 5) to MI results (Supplemental Table 6), indicates that the for both outcomes, the MI approach captures and corrects a moderate “unhappy user” attrition bias, but that the fixed time coefficients remain significant in MI analyses, indicating significant improvement in satisfaction and cramping in the six following Cu IUD placement. In all growth curve models, repeated assessment error variances were constrained equal, and random time variances were fixed, based on nested model likelihood ratio tests [15, 22].

**Table 2.** Structural equation models parameter estimates for growth curves of bleeding (PBAC Score), IUD satisfaction (5-point Likert), and Cramping (6-level MSQ item) using
multiple imputations (30 imputations)

| Outcome: | <u>Bleeding</u> | <u>IUD Satisfaction</u> | <u>Cramping</u> |
| --- | --- | --- | --- |
| Link Function: | Negative Binomial | Ordered logistic | Ordered logistic |
| Time (study month) | -0.055*<br>(-2.384) | 0.154**<br>(2.652) | -0.280***<br>(-4.378) |
| Intercept | 5.126***<br>(54.695) |  |  |
| Overdispersion (ln($\alpha$)) | -0.620***<br>(-8.039) | | |
| Random intercept variance | 0.361***<br>(4.210) | 3.458***<br>(3.839) | 4.474***<br>(4.203) |
| Threshold 1 |  | -3.529***<br>(-9.504) | -3.709***<br>(-9.743) |
| Threshold 2 |  | -2.548***<br>(-7.725) | -2.354***<br>(-6.893) |
| Threshold 3 |  | -1.087***<br>(-3.716) | 1.072***<br>(3.426) |
| Threshold 4 |  | 1.637***<br>(5.529) | 2.242***<br>(6.703) |
| Threshold 5 |  |  | 4.716***<br>(9.122) |
| N<sub>j</sub> (Assessments) | 462 | 462 | 462 |
| N<sub>i</sub> (Subjects) | 77 | 77 | 77 |

t statistics in parentheses
\* p<0.05; \*\* p<0.01; \*\*\* p<0.001
PBAC – Pictorial Blood Assessment Chart; IUD – intrauterine device; MSQ - Menstrual Symptoms Questionnaire

In our final analysis, post-insertion bleeding was significantly negatively associated with IUD satisfaction across the study (b = −1.052, p<0.05), and at study conclusion (b= −1.77, p<0.01), controlling for cramping and retrosepctive-report baseline bleeding (Table 3, Columns 2 and 3, respectively). Cramping was not significantly associated with IUD satisfaction in either specification, controlling for retrospective baseline bleeding and prospective post-placement bleeding. The sensitivity analysis shown in Supplemental Table 6 show demonstrate this finding was robust across all tested (i.e., ordered probit and linear) link functions (p<0.001). Retrospective bleeding, reported by participants at the time of enrollment, showed mixed evidence of a negative, independent effect on IUD satisfaction (Table 3: average satisfaction: p=ns; satisfaction at study end: p<0.05).

## Main Findings

New copper IUD users experience decreasing bleeding and cramping and increasing IUD satisfactionacross the first six months of Cu IUD use. Increases in overall method satisfaction across the study is significantly associated with post-insertion bleeding decreases (Table 3, Model 2: b=−1.77, p<0.01).

**Table 3:** IUD Satisfaction predicted by bleeding during study, cramping, and retrospective
report baseline bleeding; using multiple imputations (30 imputations)

| Outcome: | IUD Satisfaction<br>(study mean) | IUD Satisfaction<br>(study end) |
| --- | --- | --- |
| Link Function: | Linear | Ordered logistic |
| IUD-induced bleeding | -1.052*<br>(-2.306) | -1.766**<br>(-2.812) |
| Cramping | -0.033<br>(-0.334) | -0.084<br>(-0.752) |
| Baseline bleeding | -0.002<br>(-1.310) | -0.003*<br>(-1.963) |
| Intercept | 0.235<br>(0.913) |  |
| Threshold 1 |  | -5.003***<br>(-4.770) |
| Threshold 2 |  | -3.093***<br>(-6.047) |
| Threshold 3 |  | -1.269***<br>(-3.741) |
| Threshold 4 |  | 0.200<br>(0.648) |
| N | 77 | 77 |
| R-sq | 0.092 |  |
| Pseudo R-sq |  | 0.066 |

t statistics in parentheses
\* $p < 0.05$ ; \*\* $p < 0.01$ ; \*\*\* $p < 0.001$
IUD – intrauterine device

## Strengths

This study has a number of strengths including an adequately powered, prospective assessment, and focused analysis with high (>90%) retention rates. Additionally, the use of validated bleeding measures in combination with electronic period trackers and electronic data reporting potentially strengthen reporting reliability. Statistical methods employed in the analysis also provided a robust assessment of the impact of bleeding patterns on satisfaction. The use of SEM allowed us to avoid pooling of the means, and parse out the impact of baseline bleeding and IUD induced bleeding. The use multiple imputation allowed us to address potential attrition bias due to nonrandom missing data among women who discontinued their device due to dissatisfaction, bleeding, or cramping.

## Limitations

A main limitation was the lack of prospective pre-insertion bleeding data. We attempted to collect retrospective bleeding data leading up to IUD insertion, but felt that there were too many limitations to use this data for the primary analysis. Therefore, this manuscript focuses the findings on the prospective reports of decreasing bleeding and cramping and increasing satisfaction. Future research that incorporates prospective collection of pre-insertion bleeding using similar analytic approaches would enhance the findings presented here.

## Interpretations

Similar to other contraception studies, women report high levels of IUD method satisfaction over time [23]. New copper IUD users experience decreasing bleeding and cramping and increasing IUD satisfaction over the first six months of Cu IUD use. Overall method satisfaction across the study is associated with prospective post-insertion bleeding.

## Conclusions

Patient-centered contraceptive counseling helps individuals select methods that are best for them. These patient-driven decisions are associated with higher patient satisfaction [24]. Conversely, patients who felt their method selection was influenced by perceived provider bias towards or against a specific method reported lower satisfaction [25]. Unbiased provider counseling may help women anticipate the likelihood, duration, and intensity of side effects, provide resources for treatment plans to ameliorate side effects, and support patient′s reproductive goals. Because the most common reason for early IUD discontinuation is related to the side effects, specifically bleeding and cramping, it is important for providers to have accurate information about those side effects and the changes over time. These findings support the practical recommendation that providers give anticipatory patient counseling about their diminishing bleeding and cramping patterns. Additionally, researchers and practitioners should continue to seek practices that minimize irregular bleeding post IUD insertion as higher levels of bleeding is associated with lower satisfaction with the Cu IUD [26–28]. One option is use of a low-cost, 5-day course of over-the-counter non-steroidal anti-inflammatory drugs (NSAIDs) to decrease bleeding and cramping with menses during the first three months when most women who experience these side effects report them at the highest levels [29, 30]. Although such interventions may not improve continuation rates [29], they may accelerate the return to baseline bleeding and cramping [28]. In the absence of highly effective treatment for increased bleeding in the first several months post copper IUD insertion, medical providers should be providing women with objective information on the likely changes in bleeding they will experience. To that end, the findings from this study are helpful and reassuring.

## Acknowledgements

The authors would like to extend our deepest gratitude to Amy Orr, our exceptional research nurse and study coordinator. Additional thanks to the Consortium for Families and Health Research (CFAHR) at the University of Utah. The use of REDCap for survey and data management was provided by Eunice Kennedy Shriver National Institute of Child Health and Development grant (8UL1TR000105 (formerly UL1RR025764) NCATS/NIH).

## Supporting Information

**Supplemental Table 1. Structural equation model parameter estimates for growth curves of bleeding (PBAC Score); Unimputed, complete case analysis**

**Supplemental Table 2. Structural equation model parameter estimates for growth curves of bleeding (PBAC Score); Multiple imputation, 30 imputations**

**Supplemental Table 3. Structural equation model parameter estimates for growth curves of IUD Satisfaction (5-level Likert scale); Unimputed, complete case analysis**

**Supplemental Table 4. Structural equation model parameter estimates for growth curves of IUD Satisfaction (5-level ordinal Likert scale); Multiple imputation, 30 imputations**

**Supplemental Table 5. Structural equation model parameter estimates for growth curves of cramping (6-level ordinal MSQ item); Unimputed, complete case analysis**

**Supplemental Table 6. Structural equation model parameter estimates for growth curves of cramping (6-level ordinal MSQ item); Multiple imputation, 30 imputations**

**Supplemental Table 7. IUD Satisfaction predicted by bleeding during study and cramping; Multiple imputation, 30 imputations**

